# scGeneRythm: Using Neural Networks and Fourier Transformation to Cluster Genes by Time-Frequency Patterns in Single-Cell Data

**DOI:** 10.1101/2023.11.26.568761

**Authors:** Yiming Jia, Hao Wu, Jun Ding

## Abstract

Clustering genes in single-cell RNA sequencing plays a pivotal role in unraveling a plethora of biological processes, from cell differentiation to disease progression and metabolic pathways. Traditional time-domain methods are instrumental in certain analyses, yet they may overlook intricate relationships. For instance, genes that appear distinct in the time domain might exhibit striking similarities in the frequency domain. Recognizing this, we present scGeneRhythm, an innovative deep learning technique that employs Fourier transformation. This approach captures the rich tapestry of gene expression from both the time and frequency domains. When evaluated across a spectrum of single-cell datasets, scGeneRhythm consistently outperforms conventional approaches. The gene clusters it identifies not only demonstrate heightened statistical significance in enriched pathways but also bring to light underlying gene relationships previously obscured. Through integrating frequency-domain data, scGeneRhythm not only refines gene grouping but also uncovers pivotal biological insights, such as nuanced gene rhythmicity. By deploying scGeneRhythm, we foster a richer, multi-dimensional understanding of gene expression dynamics, enriching the potential avenues of cellular and molecular biology research.

## Introduction

Single-cell RNA sequencing (scRNA-seq) has unveiled the intricate landscape of cellular heterogeneity [1], emphasizing the need for effective gene clustering. This advanced technique has revolutionized our understanding of individual cell behaviors within complex tissues. Gene clustering in scRNA-seq analysis is paramount for deciphering this complexity, enabling insights into gene interactions, disease-associated genes [2], and the interplay between cell types [3] and biological processes [4]. Such clustering illuminates sets of genes linked with specific diseases or pathological trajectories, thereby deepening our grasp on disease mechanisms, spotlighting potential therapeutic avenues [5], and unveiling novel biomarkers for both diagnosis and prognosis [6]. Beyond pinpointing disease-associated genes, clustering also categorizes cells into distinct types or subgroups based on shared expression patterns [7]. This categorization, when integrated with data on cell morphology, function, and metabolic states, enriches our understanding of the diverse roles and relationships of cell types [8]. Furthermore, gene clustering reveals genes with synchronized expression patterns, suggesting their involvement in shared biological processes or regulatory circuits [9]. Notably, gene clustering also charts dynamic gene expression shifts during cell maturation and differentiation [10], allowing researchers to trace cellular transitions from nascent stem cell phases to their mature counterparts, and to identify the key genes and pathways orchestrating these processes.

In the burgeoning field of single-cell data analysis, the primary focus has been on categorizing cells into distinct types and modeling their developmental and differentiation trajectories [11]. However, a conspicuous gap exists in the detailed analysis of genes within these identified clusters or along specific paths of the reconstructed trajectories. Current methodologies, including scLM [12] and GPseudoClust [13], have primarily centered on gene clustering within cell types, neglecting the rich dynamic information embedded in time series scRNA-Seq studies and the pseudotime data unearthed by trajectory inference techniques. On the other hand, methods like scSTEM [14], which do consider temporal information, are primarily anchored in the time domain for gene clustering. This time-domain focus, while valuable, can miss out on capturing intricate features of gene expression data, such as low-frequency fluctuations, high-frequency fluctuations, transient gene expression behaviors, periodic patterns, oscillations and other nuanced characteristics. [15] This scenario underscores a significant need for more encompassing methods that can holistically capture the multifaceted nature of gene expression in single-cell studies.

To address the prevailing challenges in single-cell gene clustering, we present scGeneRythm, an innovative method anchored in a deep neural network framework. This approach is meticulously crafted to exploit the entirety of data within the dataset. Where traditional methods often neglect substantial temporal information, scGeneRythm ensures its holistic incorporation. Crucially, our method harnesses the frequency signal of gene expression, unveiled by the Fast Fourier Transformation (FFT), to delve deeper into gene interrelationships. Take, for example, two biologically complementary genes exhibiting contrasting expression trajectories. While time-centric methods might overlook their potential association, their frequency domain profiles could be remarkably aligned. By harmoniously integrating both time and frequency dimensions, scGeneRythm captures such intricate gene relationships with enhanced precision. Assisted by FFT, our method provides a detailed portrayal of genes, and with the power of the deep neural network, it produces gene embeddings [16] invaluable for subsequent tasks like gene visualization, clustering, and relationship quantification [17]. In extensive evaluations across diverse single-cell datasets, scGeneRythm consistently eclipsed its contemporaries, underscoring its superior gene clustering prowess. This superiority manifests in two distinct ways: firstly, through its unmatched gene clustering accuracy derived from its adept use of both time and frequency domains; and secondly, by transcending basic clustering to unearth domain features intrinsic to each gene cluster, thereby enriching them with profound biological relevance. These insights serve as a bridge, connecting raw data to meaningful biological interpretations, facilitating a more profound exploration of clustered genes’ functions and roles. scGeneRythm emerges not merely as an advanced clustering tool but as a beacon for deeper insights into the complex tapestry of gene expression in single-cell sequencing, amplifying our understanding of myriad biological processes.

## Results

### scGeneRhythm method overview

The scGeneRhythm method offers a novel approach to identifying gene sets with shared expression patterns, particularly in dynamic biological contexts such as cell growth and differentiation. A unique aspect of our method is its dual focus: while many traditional techniques emphasize the time domain, scGeneRhythm also adeptly captures the ‘rhythm’ or frequency domain of gene expression. The process begins with the construction of trajectory trees using an appropriate trajectory inference method, displayed on a UMAP [18] plot (Fig. 1). These trajectories, rooted in the time domain, are enriched with milestones serving as key nodes that interlink adjacent cells. By leveraging Fast Fourier Transform (FFT) [19], scGeneRhythm extracts frequency characteristics of these trajectories, ensuring consistent gene expression through cubic spline interpolation (Fig. 1 a). This step is crucial for identifying genes with analogous rhythmic patterns. To augment the dataset, the KNN algorithm [20] is deployed to transform the Protein-Protein Interaction (PPI) network [21] into a gene graph, making intricate connections between genes and their associations. This integration not only taps into established biological priors but also harmonizes signals from interconnected genes. A purpose-built GCN+VAE model then seamlessly amalgamates the time, rhythm, and graph data (Fig. 1 b). Once trained, this model produces concise gene embeddings that distill the inherent relationships and rhythms within the dataset. These embeddings, with their multifaceted utility, are instrumental for tasks such as gene clustering and visualization (Fig. 1 c), culminating in the discerning discovery of gene sets that share both expression patterns and rhythms.

**Fig. 1.**
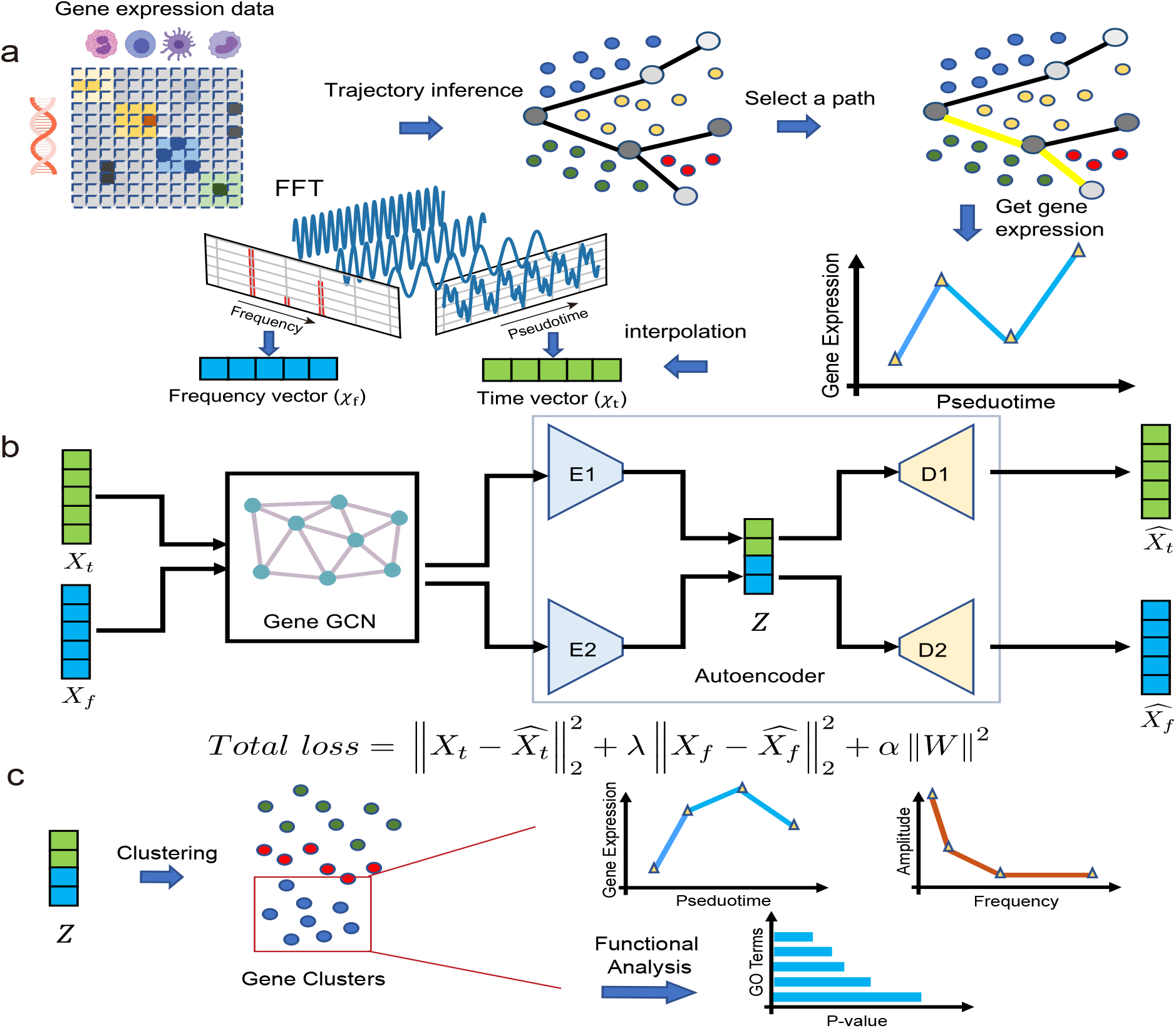
Overview of scGeneRythm. **a**, Gene expression for each waypoint is determined by averaging the expressions of all associated cells. Subsequently, a line plot is derived, representing the discrete gene expression profile in the time domain. This profile is then interpolated using spline curve fitting to obtain a continuous gene expression profile, *X*_*t*_ (marked in green). The Fast Fourier Transform (FFT) is next employed to the gene expression vector in the frequency domain, *X*_*f*_ (colored in blue). **b**, The deep neural network structure presented takes *X*_*t*_ and *X*_*f*_ as inputs. It features Graph Convolutional Networks (GCN) to leverage prior knowledge about the gene networks. Specialized encoders and decoders are incorporated to manage temporal and frequency information. This design ensures the model excels at learning gene embeddings by interpreting their expression dynamics across both the time and frequency domains. **c**, Following the acquisition of the gene embeddings, the downstream tasks encompass gene embedding clustering to identify gene clusters. The quality of these clusters is then evaluated using functional enrichment analysis, specifically through Gene Ontology (GO) term analysis. This process aids in distinguishing the unique time and frequency patterns characteristic of different gene clusters.

### scGeneRhythm effectively captures gene clusters in Mouse embryonic blood cells

In the analysis of the Mouse embryonic blood cells dataset, scGeneRythm’s effectiveness in identifying gene clusters was prominently showcased. The visual depiction in Fig. 2 a unraveled two distinct cell partitions with pseudotime paths on the UMAP plot. A closer examination of Partition 2 (highlighted in brown), abundant in primitive erythroid lineage, definitive erythroid lineage, and white blood cells, showcased a predominant path (depicted in red) selected for its thorough representation of the biological process. ScGeneRythm adeptly identified 12 gene clusters along this notable path based on the similarity of genes in both time and frequency domains, with Clusters 1 and 4 standing as testimony to the model’s precision (Fig. 2 b). A deeper examination of Fig. 2 c and d provides a comparative visualization of gene expression patterns in both time and frequency domains for Gene Cluster 1 and Gene Cluster 4, respectively. Notably, the frequency peak displayed in panel c is wider than that of panel d, indicating a broader range of rhythmic gene expression patterns inherent to Gene Cluster 1 as compared to Gene Cluster 4. This broader frequency peak may suggest that the gene expression associated with Gene Cluster 1 is more noisy and holds higher stochasticity, reflecting potentially more complex or varied regulatory mechanisms at play. The GO term enrichment analysis further substantiated the precision of scGeneRythm, unveiling significant annotations at notably low P-values. Delving into the specifics, Cluster 1 genes accentuated crucial aspects of cell and circulatory system development (e.g., ‘tube development’), while Cluster 4 genes resonated with the regulation of developmental processes like ‘regulation of anatomical structure morphogenesis’. A pivotal observation was the variance in frequency domains between the two clusters, spotlighting scGeneRythm’s adeptness in capturing underlying rhythmic patterns beyond mere temporal shifts. Illustrating the model’s analytical depth, genes GNG12 and IFNAR2 from Cluster 4 (Fig. 2 e) exhibited contrasting time domain patterns but shared frequency domain similarities. IFNAR2, a subunit of the interferon receptor essential for initiating antiviral and immune responses, alongside GNG12, integral in cell signaling as part of G-proteins potentially contributing to immune regulation, were proficiently related by scGeneRythm [22]. This insightful discernment, potentially elusive in models solely focused on temporal data, underscores scGeneRythm’s profound capability in deciphering complex gene clusters on mouse embryonic blood cells during early organogenesis. The time and frequency patterns associated with all identified gene clusters from scGeneRythm were provided in the Supplementary Figure S1.

**Fig. 2.**
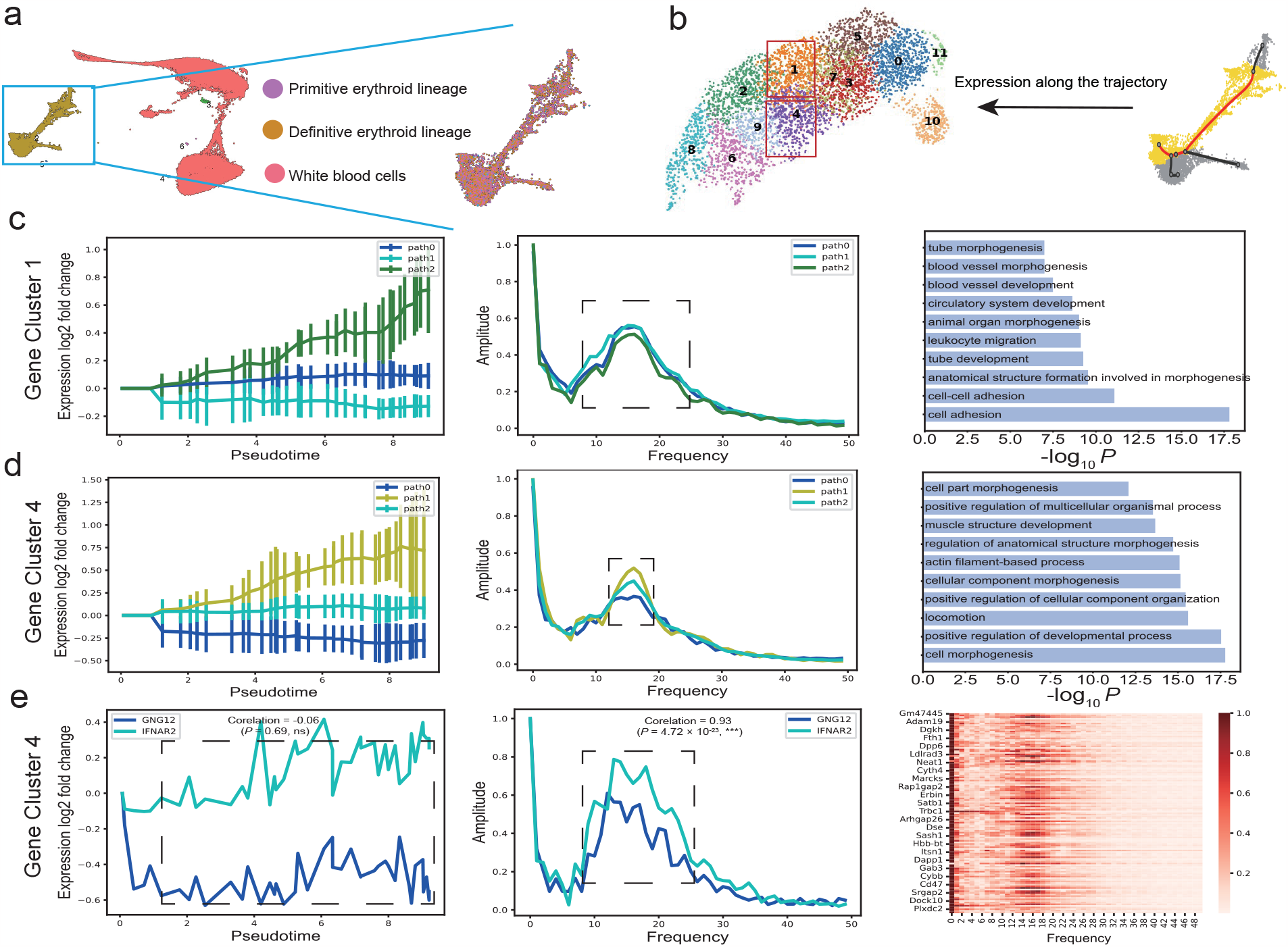
scGeneRhythm’s proficiency in delineating gene clusters within mouse embryonic blood cells dataset. **a**, UMAP visualization showcasing the Mouse embryonic blood cells dataset. **b**, Demonstrating the process of inferring a gene trajectory within the UMAP visualization using scGeneRythm, followed by clustering the resultant gene embeddings. **c**, Display of gene expression patterns for cluster 1 in both the time and frequency domains. The validity of the identified gene clusters is confirmed by the enriched Gene Ontology (GO) terms. **d**, Presentation of gene expression patterns for cluster 4, mirroring the analysis of cluster 1, and further validated by enriched GO terms. **e**, Examination of two specific genes from cluster 4, namely GNG12 and IFNAR2, revealing distinct patterns in the time domain but consistent patterns in the frequency domain. The synergistic relationship between these two genes is corroborated by existing literature. Concluding the panel is a heatmap spotlighting the frequency patterns of the top genes within cluster 4

### scGeneRhythm effectively captures gene clusters in Mouse embryonic neural crest cells

In a further exhibition of scGeneRythm’s proficiency, the algorithm was deployed on the Mouse embryonic neural crest cells dataset. This dataset predominantly captures cells instrumental for embryonic neural development, such as the Schwann cell precursor. Following processing, a singular cell partition emerged, revealing 15 pseudotime paths on a UMAP plot as depicted in Fig. 3 a. Focusing on the trajectory marked by the red line, scGeneRythm unveiled 15 distinct gene clusters, as portrayed in Fig. 3 b. These clusters adeptly pinpointed valuable gene sets, with the GO term analysis further affirming their significance through noteworthy annotations at low P-values. Transitioning to a deeper analysis, Fig.3 c-d exhibit the frequency domain dominated by a low-frequency peak centered around 0, suggesting less stochasticity in the gene expression changes in Cluster 5 and Cluster 8. This implies a more deterministic or regular expression pattern in these two gene clusters along this trajectory, showcasing scGeneRythm’s ability to capture effective gene clusters in this neural crest data. Delving into the specifics, Cluster 5 genes underscore synapse development (Fig. 3 c), featuring relevant terms like ‘synapse’ and ‘postsynaptic density’. Concurrently, Cluster 8 genes, segmented into four time domain categories (path0 to path3 as shown in Fig. 3 d), predominantly relate to morphogenesis, spotlighting terms like ‘animal organ morphogenesis’. Further exemplifying the effectiveness of scGeneRythm, genes FGF14 and CADPS in Cluster 5 are visualized in Fig. 3 e. Although these two genes exhibit distinct gene expression patterns in time domain, their frequency domain expression patterns align well. It’s pivotal to acknowledge that the genes in Cluster 5 share significant biological interconnections, particularly in neural contexts. Remarkably, both CADPS and FGF14, characterized by long introns, have been scrutinized in previous studies involving RNA samples from motor neurons in ALS patients, hinting at potential transcript reductions in these genes [23]. The analysis by scGeneRythm deftly identifies coherent gene clusters and reveals intricate relationships, even amid pronounced variations in temporal gene expression patterns. This underscores the algorithm’s prowess in harnessing both time and frequency domain features to decipher complex gene expression dynamics, especially evident in the embryonic neural crest cells dataset. The time and frequency patterns associated with all identified gene clusters from scGeneRythm for this neural creast cells dataset were provided in the Supplementary Figure S2. Beyond the above datasets, we also utilized scGeneRythm to explore human fetal immune cells using a dataset from Cao et al. [24]. As demonstrated in supplementary Figure S3 and S4, our analysis emphasized key processes like T cell activation and differentiation associated with the identified gene clusters, showcasing scGeneRythm’s prowess in deciphering crucial gene groups in fetal development and immune dynamics.

**Fig. 3.**
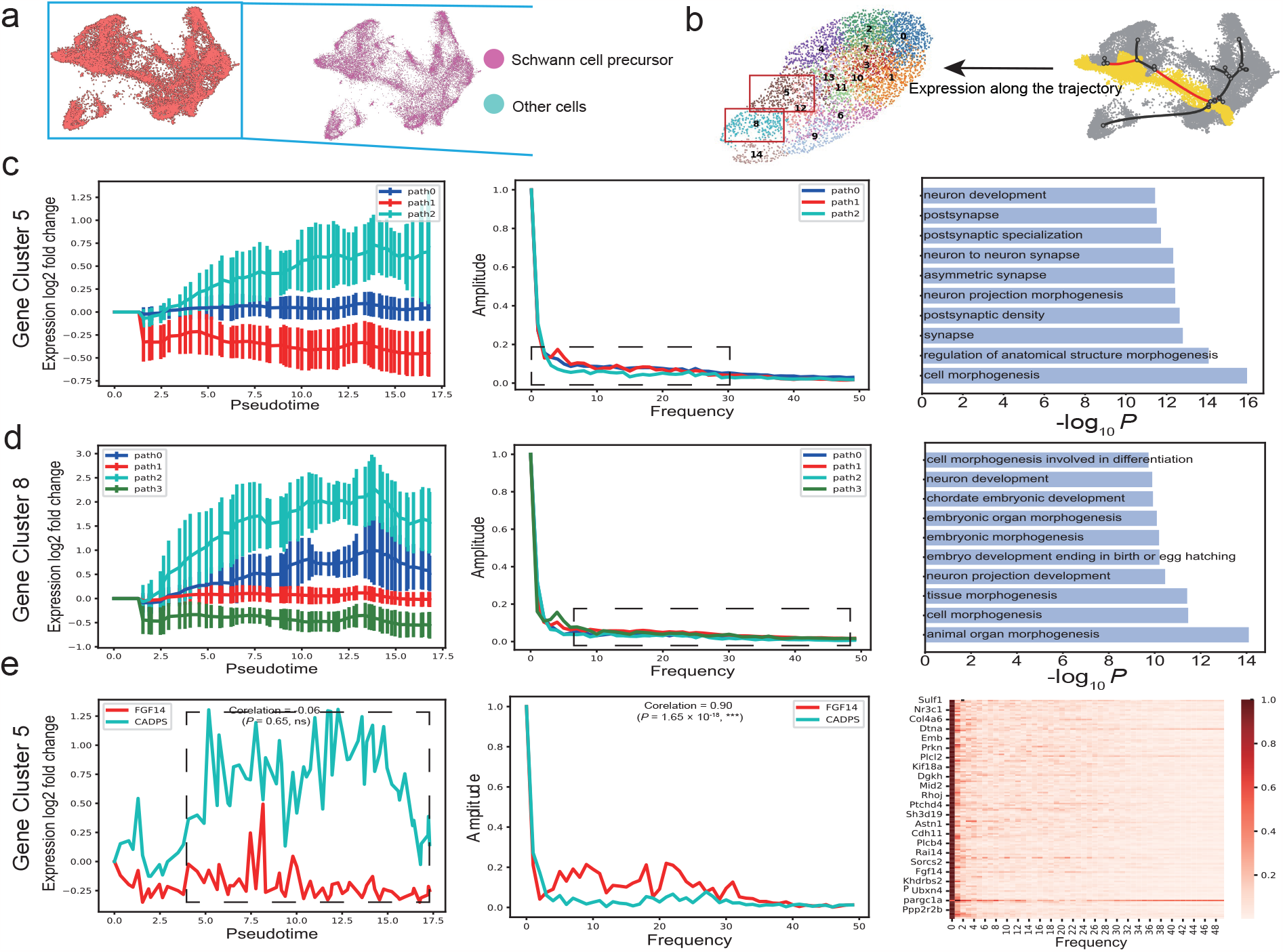
scGeneRhythm’s insights from the Mouse embryonic neural crest cells dataset. **a**, UMAP representation highlighting the Mouse embryonic neural crest cells. **b**, Depiction of the gene trajectory inferred. **c**, Analysis of gene expression patterns for cluster 5, spanning both the time and frequency domains. The efficacy of the identified gene clusters is authenticated by the enriched Gene Ontology (GO) terms. **d**, Examination of the gene expression patterns for cluster 8. **e**, FGF14 and CADPS from cluster 8 showcase distinctive patterns in the time domain but harmonized in the frequency domain. Accompanying this is a frequency domain heatmap of top genes in Cluster 5.

### Method Benchmarking

In the realm of gene clustering from time-series RNA-seq data, STEM has been recognized as a cornerstone method [25]. Advancing this lineage, scSTEM emerged tailored for single-cell data, thus positioning itself as a pertinent benchmark for modern methodologies [14]. To our best knowledge, scSTEM remains the only method for clustering genes based on their temporal changes from the time-series single-cell RNA-seq data. Given this scenario, benchmarking *scGeneRythm* against scSTEM was a methodologically sound endeavor. The Mouse embryonic blood cells dataset was employed as the testing ground for this comparative analysis. To ensure a fair comparison, both *scGeneRythm* and scSTEM were synchronized to scrutinize the identical partition and path. The comparative journey commenced with juxtaposing the Gene Ontology (GO) term analyses of example four gene clusters derived from *scGeneRythm* against all 4 clusters captured from scSTEM, as delineated in Fig. 4 a. It was discernible that *scGeneRythm*’s annotations not only clinched substantially lower P-values but also echoed more profoundly with the inherent thematic essence of the dataset. Our method yields a greater number of terms related to vascular growth, cell morphogenesis, and similar aspects that are closely aligns with the known characteristics of the data (mouse embryonic blood cells), whereas scSTEM includes a significant amount of general terms, with many terms related to neurons, which is not in line with the dataset. Delving deeper, a more nuanced GO term enrichment analysis was conducted to accentuate the comparative prowess of *scGeneRythm* over scSTEM, with the revelations depicted in Fig. 4 b. Upon aggregating the top 50 annotations from the GO term analysis for each of all clusters, the histogram and boxplot vividly portrayed *scGeneRythm*’s proclivity for garnering smaller P-values. This evidence was further solidified by a Ranksum test, which yielded a P-value of 1.06 *×* 10^*−*23^, significantly subverting the 0.05 threshold, thereby eloquently underlining the superior performance of *scGeneRythm* over scSTEM (Fig. 4 c). This benchmarking exercise unequivocally showcases *scGeneRythm*’s enhanced capability in deciphering gene clusters with a higher level of precision and thematic resonance, marking a notable stride in the ongoing evolution of gene clustering for time-series single-cell RNA-seq data.

**Fig. 4.**
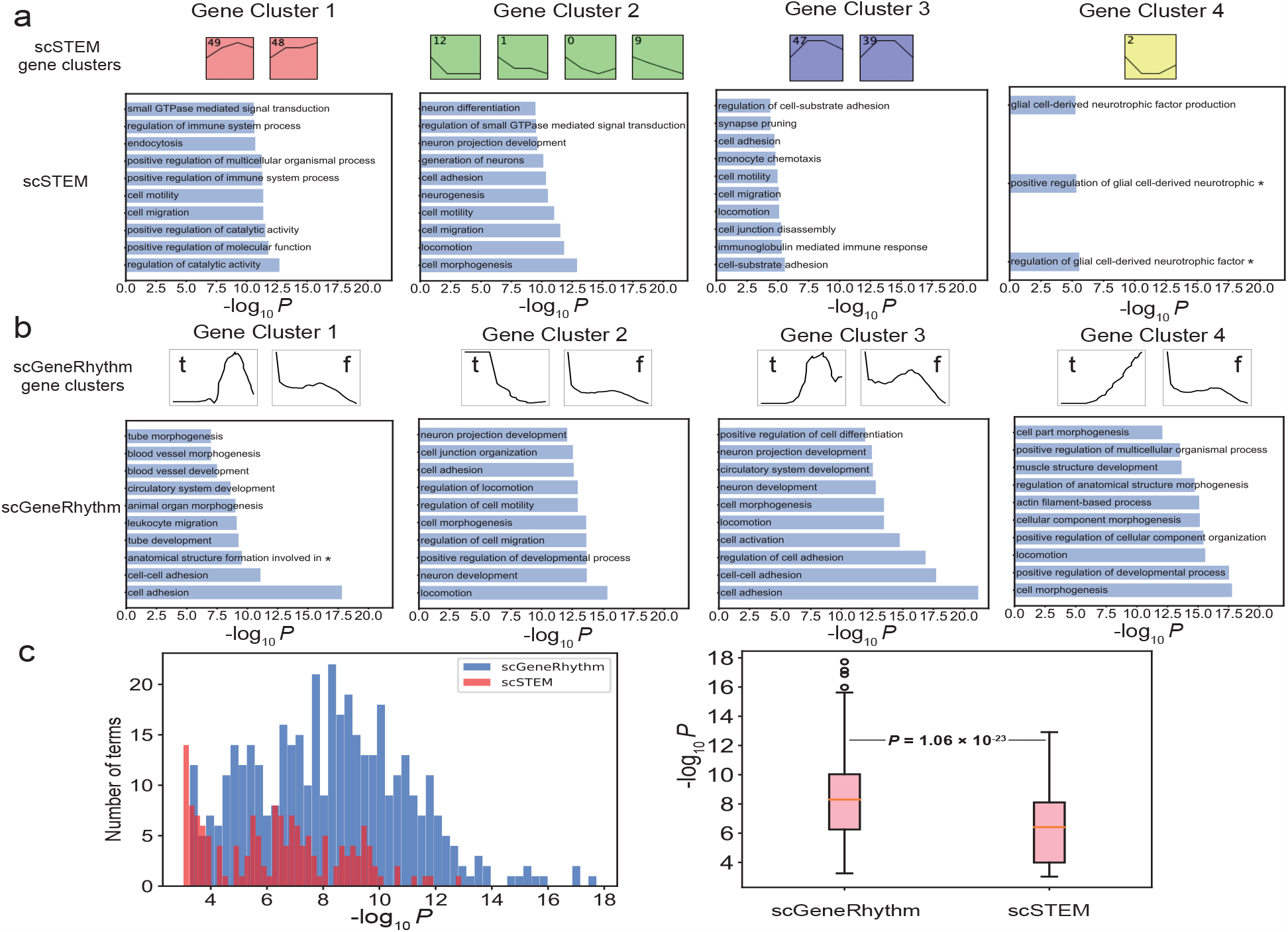
Benchmarking scGeneRythm Against scSTEM. **a**, Gene Ontology (GO) analysis for scSTEM-identified clusters, showing the top ten significant GO terms (p-values). **b**, GO analysis for top four scGeneRythm clusters, ranked by gene count, with the top ten enriched GO terms.’t’ labels indicate temporal patterns, while’f’ labels indicate frequency domain patterns. **c**, Comparative analysis using bar charts demonstrates that scGeneRythm-identified clusters exhibit more significant GO terms than scSTEM, supported by a dot plot on the right.

### Time and frequency information are both essential

To underscore the imperative of integrating both time and frequency information in *scGeneRythm*, a structured comparison was orchestrated against models singularly predicated on either time or frequency information. The ensemble of top 50 terms from the GO term analysis for each cluster was assembled, illustrated in Fig. 5 a. The comparative analysis through histograms and boxplot vividly underscored *scGeneRythm*’s heightened efficacy, thereby accentuating the pivotal role of both time and frequency features in attaining effective gene clustering. A pair of examples further epitomizes the significance of synchronizing time and frequency information, as depicted in Fig. 5 b. A model solely anchored in the time domain might erroneously link the MTSSI and SLC9A9 genes due to their similar temporal gene expression trajectories—an initial ascent followed by a decline. Yet, venturing into the frequency domain unfolds a notable divergence in their signals. The plots lucidly portray this divergence: while the peak values of both genes in the time domain are nearly synonymous, their frequency patterns markedly diverge, with peaks intersecting each other, revealing scant evidence of a substantial correlation between these genes. On the flip side, GRB10 and PID1, although showcasing limited similarity in the time domain, exhibit a pronounced resemblance in the frequency domain, as illustrated in Fig. 5 c. Biologically, both PID1 and GRB10, regulated by glucocorticoids, critically impact the insulin/IGF1 pathway; PID1 inhibits the tyrosine phosphorylation of IRS-1, while GRB10 disrupts the interaction between IR and IRS-1 [26]. This analysis robustly validates that for achieving resilient and biologically pertinent gene clustering, the fusion of both time and frequency domains is not merely beneficial, but quintessential.

**Fig. 5.**
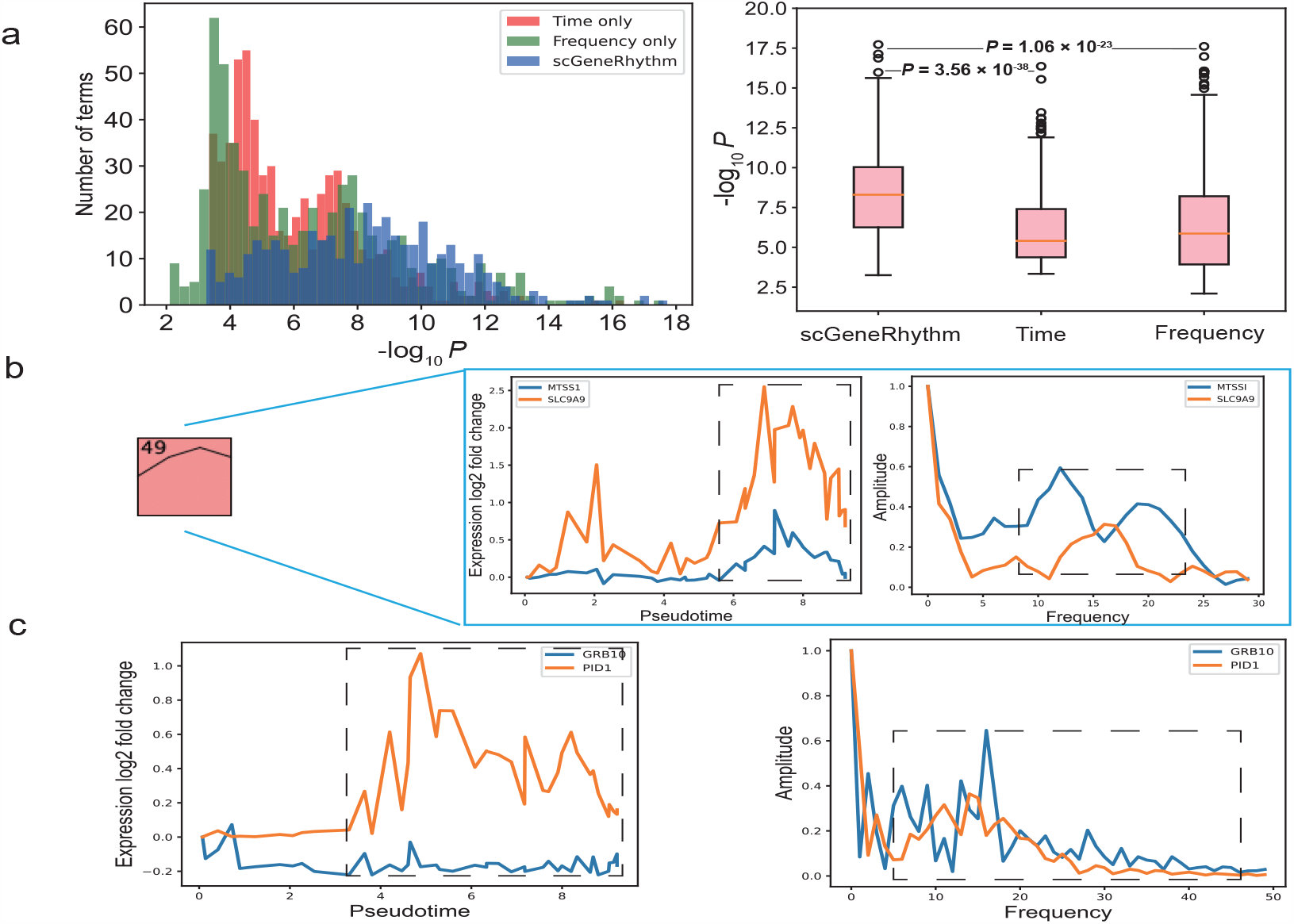
Time and frequency information are both essential. **a**, Bar chart presentation of scGeneRythm’s ablation experiments, delineating the outcomes when exclusively harnessing time domain information, solely employing frequency domain data, and utilizing our comprehensive method that synergistically integrates both domains. **b**, Through the lens of MTSS1 and SLC9A9, the limitations of an isolated time-centric perspective become manifest. Despite their concurrence in the time domain, their marked discrepancies in the frequency domain raise questions about their genuine functional collaboration. **c**, Contrasting this, the expression patterns of GRB10 and PID1 are brought to the fore, highlighting their synchronous rhythms in the frequency domain. The existing literature affirms the synergy between these genes, validating the superior efficacy of an integrated time and frequency domain approach.

## Discussion

In the presented study, we unveil scGeneRythm, a novel approach for gene clustering in single-cell RNA sequencing. Through techniques like Fast Fourier Transform (FFT) and deep learning architectures, the model fuses temporal gene expressions with their frequency counterparts, creating a unified representation.

This methodology, rigorously evaluated across a spectrum of datasets, demonstrates its prowess in pinpointing genes associated with specific diseases or pathological paths, augmenting our comprehension of disease mechanics, and spotlighting potential therapeutic strategies and biomarkers. The incorporation of Gene Ontology (GO) term analysis further underscores the model’s adeptness at extracting profound insights from complex single-cell datasets.

The brilliance of scGeneRythm emerges not only from its gene clustering capabilities but also from its revolutionary approach to conceptualizing gene expression. Central to its innovation is the pioneering introduction of gene expression frequency. By leveraging FFT, scGeneRythm navigates beyond traditional time-based genomics to unlock the frequency spectrum embedded within temporal gene expression, capturing fluctuations, rhythmic variations, and other periodicities pivotal to understanding gene behaviors. But the novelty extends further. Integrating both the time and frequency omains, our model crafts an enriched, multidimensional canvas for gene dynamics. By harmoniously weaving temporal and frequency signals, it generates a data-rich vector that becomes the bedrock for our deep learning model. This amalgamation ensures a richer, more detailed insight into gene clusters. Furthermore, the model’s architecture is laudably flexible. With the incorporation of biological priors, like the protein-protein interaction (PPI) network, scGeneRythm ensures gene clusters are not only mathematically rigorous but biologically pertinent. The modular design of our encoder and decoder architectures, bolstered by the Graph Convolutional Network (GCN) grounded in the PPI-driven gene network, empowers researchers to adapt the tool to novel genomic challenges and evolving research needs. With regards to adaptability, scGeneRythm is not only potent but incredibly versatile. Although our initial validations leaned on Seurat and Monocle, its inherent compatibility ensures seamless integration with a wide array of single-cell preprocessing and processing pipelines, aligning perfectly with tools best suited to diverse research contexts.

Through its unique integration of time and frequency gene features, scGeneRythm offers unprecedented insights into genomics. By revealing synchronized gene clusters, it surpasses traditional gene expression profiles in time domain, enabling researchers to pinpoint disruptions in disease states for improved diagnostics. Moreover, deciphering these time-frequency patterns may lead to a deeper understanding of gene expression dynamics along with disease progression via the lens of gene rhythms, potentially enhancing therapeutic efficacy. ScGeneRythm’s innovative approach has the potential to redefine how we interpret and harness the complexities of gene networks, driving genomics research to unparalleled precision and depth.

## Methods

### Single-cell RNA-seq Data Retrieval and Initial Processing

We subjected three single-cell RNA-seq datasets to scGeneRythm analysis. The first, from DESCARTES, comprises 103,766 human fetal immune cells [24]. Following the original study’s protocol, we selected the top 3,000 cell-type-specific marker genes. The other datasets, sourced from ONCOSCAPE, contained 42,262 mouse embryonic blood cells [27] and 22,283 mouse embryonic neural crest cells [27]. For these, we filtered out genes expressed in fewer than 10 cells and cells expressing fewer than 200 genes. From the refined data, we highlighted the top 5,000 most variable genes using Seurat (Version 4.0.3) [28] as our primary genes for subsequent analyses. After normalizing the single-cell cell-by-gene matrix, we condensed it to the first 100 principal components. We then employed the Leiden algorithm [29] for clustering, with the resulting clusters representing distinct cell types or unique biological processes. Detailed analysis was conducted on specific clusters, identified by their partition ID, to focus on particular cell types or processes.

### Inferring Trajectory and Pseudotime

For our chosen partition, we leveraged the monocle3 package [27] to deduce cell trajectory and pseudotime. Using the dataset’s temporal data, we identified the root node position of the trajectory and inferred each cell’s pseudotime within the partition. The trajectory was illustrated as a graph with milestone nodes and edges indicating transitions between nodes. Given that multiple paths exist from the root to leaves in the trajectory tree, each representing a potential biological process, subsequent experiments focused on individual paths, selectable by their path ID.

### Time domain feature extraction

First, we assign each cell to the milestone closest to it. Our selected trajectory path consists of several milestones, each corresponding to multiple cells. To quantify the gene expression for a milestone, we averaged the gene expressions of its associated cells. The expression for a milestone is represented as 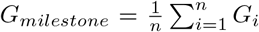, where *G*_*i*_ is the gene expression of the i-th associated cell. The pseudotime for the milestone, denoted as 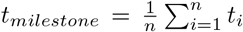, is the averaged pseudotime of these cells, with n being the total count of associated cells. Given that the milestones on the trajectory path may not be uniformly spaced in the time domain, we implemented cubic spline interpolation [30] to achieve a continuous gene expression profile. Through the aforementioned cubic spline interpolation technique, we transformed diverse sets of milestones into a standardized set of 100 pseudo time points, guaranteeing a uniform representation. As a result, each gene’s expression dynamics, mapped along its trajectory path, are captured in a 100-dimensional vector *x*_*t*_, corresponding to these consistent time points.

### Frequency Domain Feature Extraction

Once we standardized gene expression in the time domain, we then delved into understanding its frequency characteristics using the Fast Fourier Transform (FFT). FFT serves as a tool to transition a signal, often grounded in time or space, to its frequency domain counterpart, thereby revealing the periodic components that might be inherent but not immediately apparent in the original signal. To decode this, imagine the gene expression as a musical note. Just as musical notes have different frequencies determining their pitch, genes might have specific’rhythms’ or’frequencies’ to their expression. These fluctuations in the frequency domain can reflect periodic activations or inhibitions in gene expression, possibly indicating regular cellular processes or responses to external stimuli. Mathematically, the FFT in our approach is represented by:

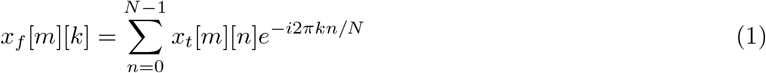

Here, *x*_*t*_[*m*][*n*] denotes the expression of the m-th gene at the n-th data point on the path, with N being the sequence’s length. The variable k varies from 0 to *N* − 1. The resulting *x*_*f*_ [*m*][*k*] is a complex number, providing insights into the amplitude and phase for the k-th frequency of the m-th gene. The amplitude is derived as |*x*_*f*_ [*m*][*k*]|. The FFT’s strength lies in its capacity to efficiently segregate gene expression into its low and high-frequency components. These frequencies can shed light on the gene’s behavior: for instance, high-frequency variations might indicate rapid changes in gene activity, while low-frequency components could represent more stable, baseline behaviors or slower responses. This conversion from 100 time domain data points translates into a concise 50-point representation in the frequency domain, offering a more distilled view of gene dynamics.

### Constructing the Gene Network Graph for Neural Network Integration

To derive richer gene embeddings within our neural network, we incorporated prior biological knowledge, notably Protein-Protein Interaction (PPI) data. We obtained PPI information from the HIPPLE database for human datasets [31] and the MIPPLE database for mouse datasets [32]. With this data related to our dataset’s genes, we employed the K-nearest neighbor (KNN) algorithm to filter out interactions with low confidence scores. This produced a refined graph structure, which serves as the foundational layer for subsequent Graph Convolutional Network (GCN) operations. By blending the neural network with this enhanced gene interaction perspective, we aim to generate gene embeddings that are both data-responsive and biologically informed.

### Deep neural network for integrating time and frequency domain features

We designed a specialized unsupervised model that synergizes the Graph Convolutional Network (GCN) [33] with the Variational Autoencoder (VAE) [34] framework. As depicted in figure 1, our model encompasses two principal modules: a GCN layer and the VAE structure which includes two encoders and two decoders. The GCN layer, informed by the protein-protein interaction (PPI) network[35], augments the relationships between genes. Subsequent to this layer, the VAE segment separately encodes time and frequency data, facilitating deeper insights from these different data modalities. These encoded representations, referred to as *Z*_*t*_ and *Z*_*f*_, are then merged and processed via a multi-layer perceptron (MLP) [36] to yield the final gene embedding, *Z*. This embedding feeds into two separate decoders for the reconstruction of the initial time and frequency data.

Formally, the time data *X*_*t*_ and frequency data *X*_*f*_ are concatenated into a 150-length tensor, which is then passed through the PPI-informed GCN layer, yielding transformed outputs 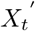 and 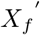. These outputs undergo encoding via *Encoder*_1_ and *Encoder*_2_ respectively. Post-encoding, the tensors *Z*_*t*_ and *Z*_*f*_ are concatenated and fed to an MLP to derive the gene embedding *Z*, which is then fully introduced to the decoders, *Decoder*_1_ and *Decoder*_2_, for reconstruction. The model’s loss function encompasses three components: reconstruction loss [37] for time data, reconstruction loss for frequency data, and a regularization term [38], articulated as:

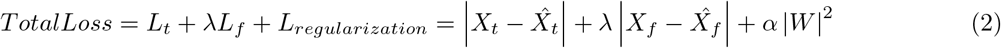

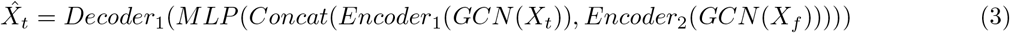

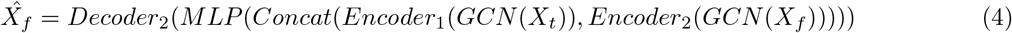

During our experimentation, we empirically determined the parameters, setting *λ* = 1 and *α* = 0. For the training phase, we utilized the AdamW optimizer [39], a prevalent algorithm known for combining momentum with adaptive learning rate strategies.

### Retrieval and clustering output gene embedding

Upon training completion, the gene expression data, encompassing both time and frequency dimensions, is fed into the encoder for inference. This produces a compact, low-dimensional representation termed as the ‘gene embedding’. Such an embedding adeptly distills and compresses the gene’s expression nuances, offering potential insights into its biological significance. Subsequently, we employ the Leiden clustering algorithm [29] on these gene embeddings, partitioning them into distinct clusters. Each cluster groups genes with congruent embeddings. Genes in the same cluster are inferred to share strong relational ties, possibly exhibiting analogous or contrasting functions, or playing pivotal roles in specific biological processes. Importantly, these clusters serve as the foundation for all subsequent analyses and examinations.

### Delineating Distinct Gene Expression Paths in time-domain within each gene cluster

While clustering the gene embeddings produced by our machine learning model enables the identification of diverse gene clusters, deeper insights can be extracted by analyzing the intricacies within each cluster. By employing k-means clustering [40] on the gene expression profiles in time-domain within a given gene cluster, we can discern distinct expression paths. These paths often exhibit pronounced disparities in the time domain, manifesting as varied trends like increasing, decreasing, etc. Interestingly, these distinctions converge in the frequency domain, rendering the paths nearly identical. This observation underscores the efficacy of our methodology in discerning gene interrelations through frequency domain data. In practice, numerous gene pairs align with this observed pattern.

### Correlation Analysis between genes in the same cluster

Our objective in utilizing correlation analysis was to examine the similarities among genes within a singular gene cluster. By assessing the correlation of gene expressions using the Pearson correlation coefficient [41], we gauged how genes are related in both the time and frequency domains. This approach provided insights into whether genes in the same cluster exhibited similarities in their expression dynamics over time (time domain) or in their oscillation patterns (frequency domain). This analytical lens enabled us to understand the nuanced relationships and dynamics between genes within a cluster more deeply.

### Gene Cluster Evaluation and Benchmarking

Gene Ontology (GO) Enrichment Analysis [42] offers a structured approach to identify gene sets overrepresented in particular functional annotations. In the context of scGeneRythm, this method plays an integral role in unveiling the biological roles and related annotations for genes present in each cluster. For the enrichment analysis, we have opted for Toppfun [43], which stands out as a holistic tool tailored for enrichment analysis [44] of gene lists and also offers capabilities for candidate gene prioritization, drawing on functional annotations and protein interaction networks. For our analyses, we feed gene lists, curated from each cluster via the Leiden algorithm, directly into Toppfun, from which we extract enriched GO terms. Our analysis particularly hones in on the annotations cataloged under the GO: Biological Process category, providing insights into the biological undertakings associated with the listed genes. When assessing individual gene clusters, our attention is drawn to the most significant GO terms from the Biological Process category, particularly those with the smallest p-values, all the while ensuring they surpass a p-value benchmark of 0.05. With the rank-sum test [45] in our analytical toolkit, we undertake a broad comparison across gene cluster distributions, facilitating an understanding of the validity and nuanced differences in GO terms derived from an assortment of methods.

## Software availability

scGeneRythm is a tool designed for single-cell level gene clustering, leveraging both time and frequency domain information. It is open-sourced on GitHub at https://github.com/jymmmmm/GeneRythm, complete with detailed instructions, documentation, and examples.

## Acknowledgements

This work was funded in part by grants awarded to [JD]. We gratefully acknowledge the support from the Canadian Institutes of Health Research (CIHR) under Grant Nos. PJT-180505; the Funds de recherche du Québec - Santé (FRQS) under Grant Nos. 295298 and 295299; the Natural Sciences and Engineering Research Council of Canada (NSERC) under Grant No. RGPIN2022-04399;; and the Meakins-Christie Chair in Respiratory Research. The work is also funded in part by grants award to [HW] from the National Natural Science Foundation of China (NSFC) under Grant Nos. 62272278 and 61972322.

